# Visual evoked potentials as an early-stage biomarker in the rTg4510 tauopathy mouse model

**DOI:** 10.1101/2022.10.03.510063

**Authors:** Aleksandra Parka, Christiane Volbracht, Benjamin Hall, Jesper F. Bastlund, Maiken Nedergaard, Bettina Laursen, Paolo Botta, Florence Sotty

## Abstract

Tauopathies such as Alzheimer’s Disease (AD) and frontotemporal dementia (FTD) are characterized by formation of neurofibrillary tangles consisting of hyperphosphorylated tau protein. Early pathophysiological and functional changes related to neurofibrillary tangles formation are considered to occur prior to extensive neurodegeneration. Hyperphosphorylated tau has been detected in postmortem retinas of AD and FTD patients, and the visual pathway is an easily accessible system in a clinical setting. Hence, assessment of the visual function may offer the potential to detect consequences of early tau pathology in patients. In this study we explored the association between the visual system and functional consequences of tau pathology progression using a tauopathy rTg4510 mouse model. To this end, we recorded full-field electroretinography and visual evoked potentials in anesthetized and awake states at different ages. While retinal function remained mostly intact within all the age groups investigated, we detected significant changes in amplitudes of visual evoked potential responses in young rTg4510 mice exhibiting early tau pathology prior to neurodegeneration. These functional alterations in the visual cortex were positively correlated with pathological tau levels. Our findings suggest that visual processing could be useful as a novel electrophysiological biomarker for early stages of tauopathy.

## Introduction

Tauopathies are neurodegenerative diseases characterized by pathological tau hyperphosphorylation and accumulation. The most common tauopathy is Alzheimer’s Disease (AD) that is currently affecting 55 million people and ranking as 7^th^ leading cause of mortality worldwide [1]. Tau is a microtubule associated protein encoded by the MAPT gene [2], which under physiological conditions, among other functions, plays an intrinsic role to stabilize microtubules and is involved in axonal transport [3]. Disruptions to normal tau functions by hyperphosphorylation, oligomerization, and aggregation occur in tauopathies and are considered to obstruct neuronal transport, finally leading to neuronal death [4]. Together with amyloid-beta (Aβ) plaques, neurofibrillary tangles consisting of hyperphosphorylated tau are pathological hallmarks of AD [5]. Studies showed that while Aβ and tau pathologies are inherently associated, the progression of neurofibrillary tangles is closely correlated with cognitive decline in AD [6,7]. In other tauopathies such as frontotemporal dementia (FTD) neurofibrillary tangles are the primary pathological hallmark [2]. Tau hyperphosphorylation, oligomerization and aggregation is a gradual process and the initial pathophysiology occurs years prior to emergence of cognitive symptoms, which are related to irreversible brain atrophy [8].

Beside pathological findings in the brain, retinal thinning and hyperphosphorylated tau were detected in postmortem retinas of FTD and AD patients [9–11] as well as in rodent tauopathy models [12–14]. Moreover, alterations in visual evoked potentials (VEPs) have been identified in patients with high risk of developing AD [15]. Functional impairments in cortical visual plasticity and retinal responses were also detected in animal models overexpressing human tau [14,16–18]. As electroretinography (ERG) and VEP measurements are cost-effective and non-invasive methods to measure basic neuronal functioning as compared to current diagnostic tools [1], they could be employed to detect early signs of tauopathies.

Tau transgenic mice, rTg4510, overexpress the human 4R0N isoform tau with P301L mutation found in a familial form of FTD, under the calcium/calmodulin dependent kinase II alpha (CaMKIIα) promoter [19]. The resulting human tau overexpression leads to progressive tau pathology mainly in pyramidal neurons of the forebrain [20,21]. Neurodegeneration in these mice was detected in the hippocampal CA1 around 6-12 months of age and was preceded by cognitive decline as portrayed by deficiencies in memory oriented behavioral tests and by non-mnemonic behavioral defects [19,21–23]. Studies investigating early tau pathology in rTg4510 found indications of synaptic loss, changes in pyramidal neuron function and impaired neuronal activity in cortical and hippocampal areas [22,24–27]. While most rodent studies have focused on hippocampal areas in relation to their involvement in cognitive function, cortical areas are easier to study electrophysiologically in humans compared to subcortical structures. Since endogenous human tau is expressed both in the retina and visual cortex, evaluation of visual processing could represent a potential alternative functional biomarker readout of tau pathology in patients. rTg4510 mice are a suitable model to investigate effects of tau pathology on the visual system as human tau was detected in both retina and visual cortex in these mice [14,17].

The aim of this study was to assess whether early signs of hyperphosphorylated tau related pathology could be detected using a combination of *in vivo* electrophysiological recordings, immunoblotting and immunohistochemistry from the retina and primary visual cortex.

## Materials and methods

### Animals

rTg4510, tTA and non-transgenic littermate F1 mice were bred at Taconic, Denmark. Male mice were exclusively used in this study to avoid potential variability caused by sex differences [28]. Mice expressing the tetracycline transactivator transgene (tTA) under CaMKIIα promoter were maintained on 129S6 background strain (Taconic) and tau_P301L_ responder mice containing the tetracycline operon-responsive element were maintained in the FVB/NCrl background strain (Taconic). Bitransgenic animals - rTg4510 - produce the investigated tauopathy phenotype [19]. In this study, tTA littermates were used as controls (Fig. S1). Upon arrival, mice were single housed in cages containing environmental enrichment.

All animals had *ad libitum* access to water and food (Brogaarden, Denmark). The light/dark cycle was maintained at 12h, room temperature was 21 ± 2 C and a relative humidity of 55% ± 5%. The animal experiments were performed in accordance with the European Communities Council Directive no. 86/609, the directives of the Danish National Committee on Animal Research Ethics, and Danish legislation on experimental animals (license no. 2014-15-0201-00339).

### Stereotaxic surgery for visual evoked potentials recordings

All tools and surfaces were wiped with 70% ethanol prior to surgery. Mice were anesthetized with isoflurane (30% oxygen/70% nitrogen; 5% isoflurane for induction, 1.5-2% for maintenance). The scalp was shaved and disinfected with chlorhexidine followed by a subcutaneous injection of Marcain (2.5 mg/ml bupivacaine, AstraZeneca, Albertslund, Denmark). A vaseline-based gel was applied to the eyes to avoid drying (Neutral Ophtha). An incision was made along the midline and a spiral drill tip (HSS 203 008, Meisinger, Germany) was used to drill holes for screw electrodes (E363/20/1.6/SPC, PlasticsOne, Roanoke, VA, USA). For the VEP recordings in anesthetized animals, a hole was drilled above the right posterior visual cortex at the following coordinates (AP: −3.6 mm, ML: 2.3 mm) according to the Franklin & Paxinos mouse brain atlas [29]. The reference electrode was inserted in the frontal part of the skull contralaterally to the recording electrode. All screws were covered with Fuji plus cement (GC America, US) leaving connectors exposed. The mice were weighed and treated with Noromox (150 mg/ml amoxycilin hydrate, ScanVet, Fredernsborg, Denmark) and Norodyl (50 mg/ml carpofen, ScanVet, Fredensborg, Denmark) for 5 days post-surgery, and after minimum 7 days from surgery, the animals were submitted to VEP recordings.

For awake VEP recordings, coordinates for the visual cortex and screw electrodes were the same as for the animal recordings performed during anesthesia. Five additional electrodes were installed with the purpose of stabilizing the implant. The recordings from these electrodes were not used. Connectors were grouped together in a pedestal (MS363, PlasticsOne) and cemented at the base with RelyX™ Unicem dental cement (3M, Copenhagen, Denmark) and further covered with Fuji plus cement (GC America, US). Post-operative care protocol and recovery period were the same as for the surgery prior to the recordings in anaesthetized mice.

### Electroretinography and visual evoked potentials recordings during anesthesia

Scotopic ERG and VEP recordings were performed in rTg4510 and tTA (control) mice at different ages (3, 6, 9 and 16 months). Each animal was dark adapted for at least 8h before the recording session. Full-field ERG recordings were carried out using Celeris set up and data was acquired with the Espion system (both from Diagnosys LLC, Cambridge, UK). The room was maintained in complete darkness with the only red light source provided by the overhead lamp from the Celeris system.

The animals were weighed then sedated with 10 ml/kg i.p. of ketamine and xylazine cocktail (80/8 mg/kg, respectively). The pupils were fully dilated with drops containing 0.2 mg/ml tropicamide, 3.1 mg/ml phenylephrine hydrochloride and 10 mg/ml lidocaine hydrochloride (Mydrane, Thea Nordic, Hørsholm, Denmark). After 3-5 min, the excess drops were removed and 2.5% hypromellose gel was applied to the cornea to maintain the moisture and conductivity between the eye and electrode. The recording electrodes were gently placed upon the corneas. A ground electrode was inserted in the base of the tail and the VEP and reference electrodes were surgically pre-inserted in the skull. The overhead lamp was turned off to begin the recording 10 minutes later to ensure complete dark adaptation.

The signal was sampled at 2 kHz and the flash VEP stimulation protocol consisted of white flashes each lasting 14 ms. Stimulus was presented in incrementing intensities (0.001, 0.003, 0.01, 0.03, 0.1, 0.3, 1,10 cd*s/m^2^), and each intensity was repeated 30 times except for 1 & 10 cd*s/m^2^ which were repeated 15 and 10 times, respectively. The interstimulus interval was 5 s for four of the dimmest stimuli, and 7, 8, 10 and 20 s for 0.1, 0.3, 1 and 10 cd*s/m^2^, respectively. During the recordings, the eyes were interchangeably stimulated to avoid potential transfer of light between the retinas.

Photopic negative response (PhNR) recordings were performed in mice adapted to light for at least 10 minutes. A white 14 ms flash of light at 22.76 cd*s/m^2^ on a green background at 40 cd*s/m^2^ was presented 100 times at 2 Hz.

### Awake visual evoked potentials recordings

Recordings took place during the dark phase in a chamber with a light panel suspended above the cage containing nesting material. Following recovery from surgery, 6 month old animals were habituated to the recording chambers for 3h once a week for 2 weeks prior to the recording session. Whole-field light flashes were generated by LED panels suspended approximately 40 cm above the base of the cage. White light stimuli were presented for 10 ms with an inter-stimulus interval of 3 s for 0.8 and 3.44 cd*s/m^2^ (150 repeats), 4 s for 7.2 cd*s/m^2^ (150 repeats), 5 s for 11.8 and 17.3 cd*s/m^2^ (both 100 repeats), 7 s for 22.4 cd*s/m^2^ (75 repeats) and 10 s for 28.3 cd*s/m^2^ (50 repeats). The signal was amplified 1000x and a band-pass filter between 0.01-300 Hz was applied (Precision Model 440; Brownlee, Palo Alto, CA, USA). Sampling was done at 1 kHz (CED Power 1401, Cambridge Electronic Design Ltd, Cambridge, UK). Stimulation triggers were generated by a custom-made configuration in Spike2 version 7.2 (Cambridge Electronic Design Ltd, Cambridge, UK).

### Data analysis

Output of the anaesthetized ERG/VEP recordings was analyzed in MATLAB (MathWorks, Natick, Massachusetts, USA). ERG recordings were averaged between right and left eye stimulation readouts. The a-wave is the first negative deflection of the ERG waveform and is presented as the amplitude of the trough from baseline, while the b-wave corresponds to the peak amplitude of the following positive peak and is relative to the a-wave (Fig. 1A). In the VEP waveforms obtained from anaesthetized mice, N1 was measured as the amplitude of the first negative peak after the stimulus relative to baseline while P2 was measured as the amplitude of the subsequent positive peak relative to baseline. In the VEP recordings of awake mice, two additional components were identified: N2 reflecting the amplitude of the following negative deflection as well as P3 which was identified as the positive peak response of the period from 0.3-1 s post-stimulus (both components relative to baseline) (Fig 2A & 3A). Amplitudes and latencies of each component were compared between age-matched controls and rTg4510 animals. The recording electrode was implanted in the right hemisphere. Thus, due to the contralateral nature of visual pathway signaling, only the results from stimulation of the left eye were considered.

**Figure 1.**
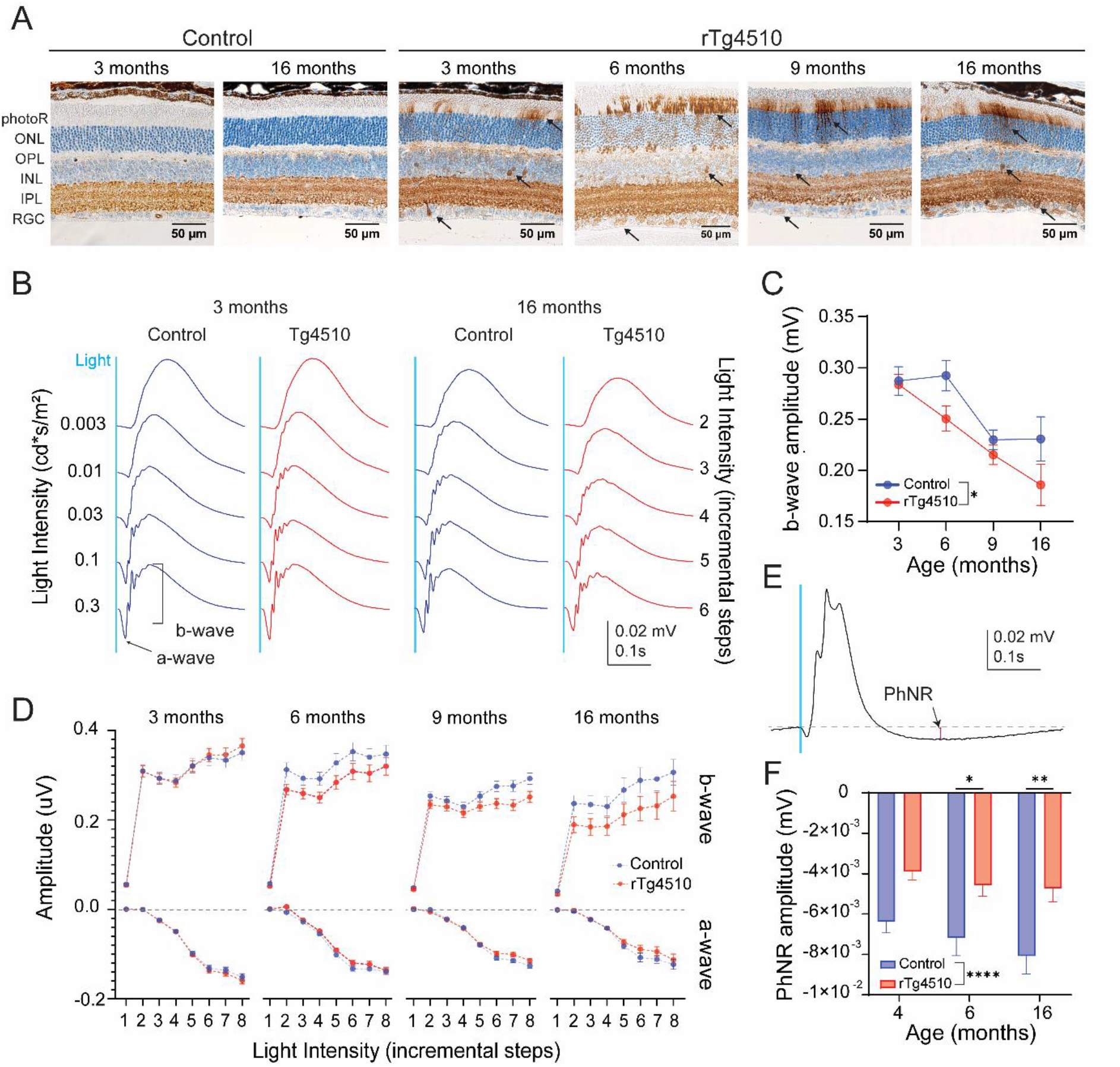
Retinal tau pathology and impact on ERG responses in control and rTg4510 mice. (A) Representative AT8 immunostainings of the retinas of control and rTg4510 at different ages. Black arrows point to exemplary abnormal AT8+ staining. Data are shown as mean ± SEM. (B) Representative waveforms of retinal responses to a white flash stimulus at various intensities in 3 and 16 month old control and rTg4510 mice. Blue line indicates the timing of the light stimulus. (C) Amplitudes of the b-wave evoked by 0.1 cd*sm^2^ of light in control (blue, n=15-22) and rTg4510 (red, n=19-24) mice at 3, 6, 9 and 16 months of age. Two-way ANOVA, (*) p<0.05. (D) Intensity-dependent amplitudes of a- and b-waves in control (blue, n=15-22) and rTg4510 (red, n=19-24) mice at 3, 6, 9, and 16 months of age. X-axis represents stimulation intensities (step 1-8 for 0.003, 0.001, 0.03, 0.01, 0.3, 0.1, 1, 10 cd*sm^2^, respectively) (E) Representative trace of PhNR. Red line indicates the amplitude of PhNR. (F) PhNR amplitudes at 4, 6 and 16 month old control and rTg4510 mice (n=10-16, each data point is represented by the readout from one eye). Two-way ANOVA (*) p<0.05, (**) p<0.01 (****) p<0.0001.

**Figure 2.**
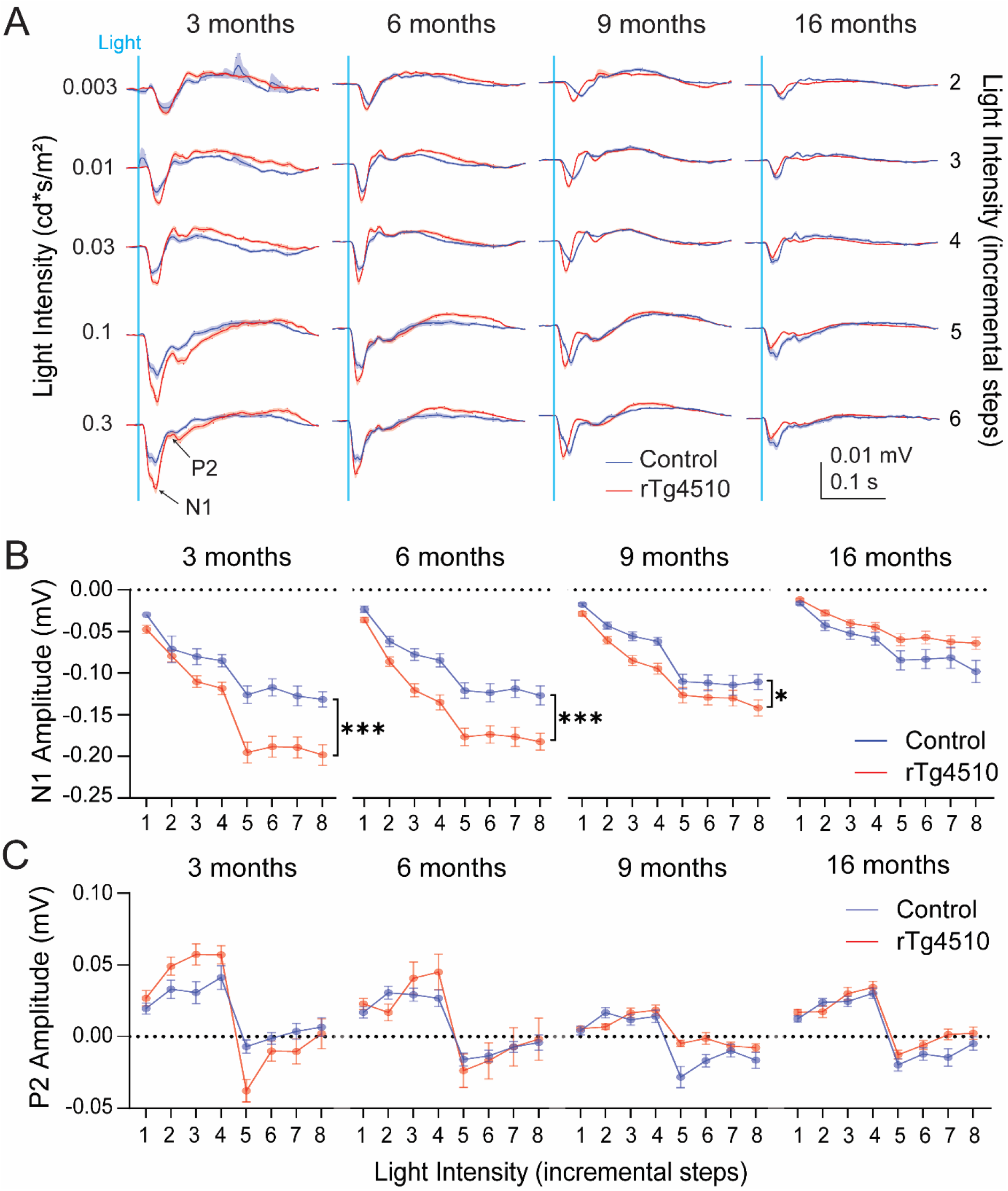
VEP recordings from anaesthetized control and rTg4510 mice responding to a white light stimulus. (A) Mean waveforms of the VEP responses from control (blue, n=15-22) and rTg4510 (red, n=19-24) mice in all age groups and different light intensities. Shaded areas along the waveforms represent SEM. (B) Average amplitudes of the N1 and (C) P2 component of the VEP in anaesthetized control (blue, n=15-22) and rTg4510 (red, n=19-24) mice at 3, 6, 9, and 16 months of age at increasing intensities. X-axis represents stimulation intensities (1-8 for 0.003, 0.001, 0.03, 0.01, 0.3, 0.1, 1, 10 cd*sm^2^, respectively). Two-way ANOVA, (*) p<0.05, (***) p<0.001. Data are shown as mean ± SEM.

**Figure 3.**
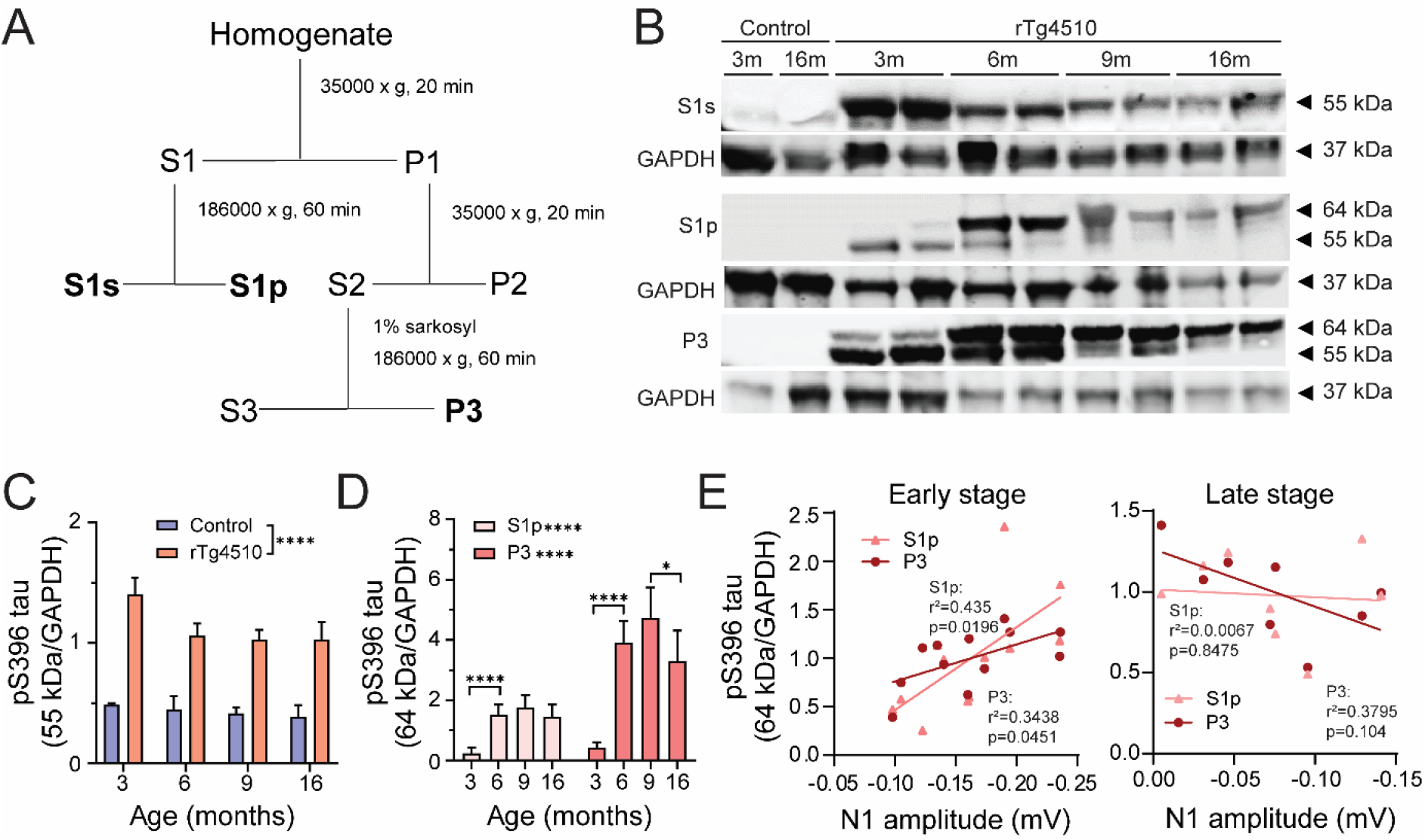
Progression of tau hyperphosphorylation in rTg4510 mice. (A) Tau fractionation protocol. S1s contains 55 kDa tau representing the levels of normal phosphorylated tau. S1p fraction contains 64 kDa tau representing soluble oligomeric hyperphosphorylated tau. P3 fraction consists of 64 kDa tau indicative of filamentous aggregated sarkosyl insoluble tau. Adapted from Sahara et. al [30]. (B) Representative gels from control and rTg4510 brain homogenates at 3, 6, 9, and 16 months of age. Fractions S1s, S1p and P3 were probed with an antibody targeting tau phosphorylated at S396 epitope. For normalization membranes were re-probed for GAPDH (C) Quantification of 55 kDa in control (blue, n=3) and rTg4510 (red, n=5-6) from S1s fraction and (D) 64 kDa tau from S1s and P3 fractions. Two-way ANOVA, S1s: genotype effect, S1s and P3: age effect (*) p<0.05, (**) p<0.01, (***) p<0.001. (E) Correlation of N1 of anesthetized VEP and S1p and P3 hyperphosphorylated tau in early (3-6 month old rTg4510; left) and late (9-16 month old rTg4510; right) stage of pathology. Data are shown as mean ± SEM.

### Tissue collection

Immediately after each ERG/VEP recording in anesthetized animals, the mice were transcardially perfused with 0.3% heparinized (10000 units/L) 0.1 M potassium phosphate buffered saline (KPBS). The eyes were enucleated and kept in Davidson’s (or Hartmann’s) fixative for 48h and then transferred to 0.01% sodium azide in 0.1 M KPBS solution. In addition, the right hemisphere of the brain was drop fixed in 4% paraformaldehyde for 48h and then transferred to 0.01% sodium azide in 0.1 M KPBS solution until histological processing. The left half of the brain was snap frozen and kept at −80°C until homogenization for Western Blot (WB) analysis.

### Immunohistochemistry

The half-brains and eyes were embedded in paraffin blocks and sectioned on a microtome into 4 μm sections. 4-5 sections across the visual cortex approximately 32μm apart from 4-5 animals per group were used. First, the sections were kept at 60°C for 30 min followed by deparaffinization in xylene. Antigen retrieval was done in PT Link Pre-treatment Module (DAKO, Glostrup, Denmark) where the sections were heated up to 95°C in citrate buffer. After cool-down, the slides were processed in Autostainer Link 48 (DAKO, Glostrup, Denmark) and incubated with AT8 (1:1000, #MN1020, Invitrogen, USA) primary antibody against pS202/pT205 tau followed by horseradish peroxidase (HRP) conjugated secondary antibody. The staining was visualized with diaminobenzidine (DAB). AT8+ cells were counted in the visual cortex and ventral hippocampus using ImageJ software (National Institutes of Health, USA), normalized to counting area and averaged between all sections from one animal. Cortical thickness was calculated as the average of 3 measurements between the top of the cortex to corpus callosum using ImageJ.

### Fractionation protocol for Western Blotting

The cerebellum was removed from the hemispheres collected for tau detection. Fractionation protocol was adapted from Sahara et. al and Helboe et. al [22,30]. Briefly, homogenized tissues were centrifuged at 35000 x g for 20 min at 4 °C resulting in supernatant (S1) and pellet (P2) fractions. The S1 fraction was further centrifuged in 186000 x g for 60 min at 4 °C. The pellet from this centrifugation was labelled S1p and resuspended in Tris-buffered saline and the supernatant S1s. P2 fraction was re-homogenized in high salt/sucrose buffer. After centrifugation in 35000 x g for 20 min at 4 °C, the supernatant was incubated with 1% sarkosyl (v/v) (Sigma, Denmark) at 37 °C for 60 min. Finally, the supernatant was centrifuged at 186000 x g for 60 min at 4 °C, and the pellet (P3) was resuspended in 0.5 volumes of TE buffer. Fractionation protocol was performed to isolate tau in various levels of phosphorylation and oligomerization. Fraction S1p contains sarkosyl soluble pre-tangle equivalent hyperphosphorylated tau while P3 represents sarkosyl insoluble fibrillar tangle species [19,30].

### Western Blotting

S1s, S1p and P3 samples containing buffer with 0.1 DTT were treated at 95 °C for 10 min and separated in a 15-well comb 4-12% Bis-Tris SDS-PAGE gel. Afterwards, the gels were transferred to a Immobilon®-FL membranes (PVDF-FL, MerckMillipore, Germany) which were later blocked in Odyssey® Blocking buffer and incubated with primary antibodies overnight at 4°C. Phosphorylated tau epitopes were stained with a rabbit pS396 monoclonal antibody (1:1000, #44-752G, Invitrogen, USA) and as a loading control, anti-GAPDH mouse antibody was used (1:1000, #MAB374, Invitrogen, USA). After the primary antibody incubation, the membranes were washed with TBST and moved to secondary antibody incubation with 800 nm anti-mouse (1:10000, #926-3221, Li-cor BioSciences, USA) and a 680 nm anti-rabbit (1:20000, #926-68073, Li-cor BioSciences, USA). Finally, the membranes were visualized in Odyssey® CLx scanner (Li-cor BioSciences, USA). The pS396 signal was normalized to that of GAPDH to obtain the level of phosphorylated tau fractions.

### Statistics

Statistics were performed in GraphPad Prism software (GraphPad Software, San Diego, California, USA). Two-way ANOVA was used when analyzing data between two age-matched genotypes at multiple stimulation intensities. If a significant effect of the genotype was detected, a Šídák’s multiple comparison post hoc analysis was performed. One-way ANOVA was employed when comparing data within genotype at different ages. Simple linear regression analysis was performed when testing correlations between two parameters. In comparisons between means of two groups, two-tailed student t-test was employed.

## Results

### Non-transgenic and tTA littermates display similar electrophysiological profile

Bitransgenic rTg4510 mice exhibit human tau overexpression driven by the calcium/calmodulin dependent kinase II (CaMKII) promoter under control of the tetracycline transactivator (tTA) [19]. The tTA transgene has previously been linked to brain changes affecting the phenotype [22,31,32]. Hence, we first investigated whether the tTA transgene could lead to differences in the electrophysiological profile in tTA compared to non-transgenic littermates. We measured no significant differences between the two genotypes at 3, 6, and 9 months of age (Fig. S1). Nonetheless, based on previous reports [22,31], we considered the tTA as the most appropriate control for rTg4510 mice and therefore, employed only tTA littermates as control animals in the subsequent experiments.

### Retinal responses decline with age in rTg4510 mice

In the retina, the progression of tau pathology in rTg4510 mice was characterized by tau phosphorylation at pS202/T205 epitopes by AT8-immunoreactivity. AT8 immunoreactivity was detected in the photoreceptor (photoR) layer, inner nuclear layer (INL) and retinal ganglion cell (RGC) layer at 3 month and consistently in 6, 9 and 16 month old rTg4510, while it was absent in control mice (Fig. 1A). There was unspecific AT8 antibody binding in the inner plexiform layer (IPL) as the same immunoreactivity was found in both control and rTg4510 animals.

To further assess the impact of tau pathology on retinal function in rTg4510 mice, we conducted scotopic ERG recordings to evaluate rod photoR function, rod-mediated bipolar cell responses and RGC function. Light stimulation elicited a clear ERG response consisting of a- and b-waves (Fig. 1B). The amplitude of the a-wave as a function of the stimulation intensity was not significantly different between rTg4510 and tTA littermates at any of the ages tested (Fig. 1D). At the intensity of 0.1 cd*s/m^2^ which evoked maximal ERG amplitude, a-wave amplitude significantly declined with age in both rTg4510 and control mice (One-way ANOVA, p = 0.0001, 0.0172 for control and rTg4510, respectively; Fig. S2). The amplitude of the b-wave as a function of stimulation intensity in rTg4510 tended to be lower compared to controls from 6 months of age, although it did not reach significance (Two-way ANOVA, p = 0.088, 0.059, 0.12 for 6, 9 and 16 months, respectively; Fig 1D). In addition, the b-wave amplitude triggered by 0.1 cd*s/m^2^ significantly declined with age in both groups (One-way ANOVA, p=0.0017 and <0.0001 for control and rTg4510, respectively; Fig. 1C), although a more severe decline was observed in rTg4510 mice compared to control mice (Two-way ANOVA, p=0.02; Fig. 1C). Latencies of a- and b-waves were not significantly different between genotypes (Fig. S3).

As retinal ganglion cells do not contribute to the full-field flash ERG response, PhNR recordings were carried out (Fig. 1E). Overall, the PhNR amplitude was significantly different between genotypes (Two-way ANOVA, p<0.0001; Fig. 1F). Post-hoc analysis showed a significant reduction in PhNR amplitude in 6 and 16 month old rTg4510 mice (Two-way ANOVA, p=0.03 and p=0.004 for 6 and 16 months, respectively).

In summary, an age dependent decline in a-wave recordings was observed in both control and rTg4510 mice, and is thereby independent of tau pathology. On the other hand, the decline in b-wave and PhNR recordings was associated with the progression of tau pathology in the retina of old rTg4510 mice.

### Enhanced amplitudes of the N1 of the visual evoked potential response in young rTg4510

After finding defects in retina function in old rTg4510 mice, we investigated the visual cortex functionality as the visual processing center. In combination with ERG recordings, we monitored cortical VEP responses during the same light stimulation protocol (Fig. 2A). Amplitudes of N1, but not P2, increased with incremental light intensity in both rTg4510 and control mice (Two-way ANOVA, p<0.0001 for 3, 6, 9 and 16 months; Fig. 2B-C). Interestingly, the amplitude of the N1 component of the VEP was significantly enhanced for nearly all intensities in 3, 6 and 9 month old rTg4510 compared to age-matched control animals (Two-way ANOVA, p=0.0005, p=0.0004 and p=0.027 for 3, 6 and 9 months, respectively; Fig. 2B). The increased N1 amplitude in rTg4510 was most prominent in 3 and 6 month old mice. In 16 month old rTg4510, there was a trend towards decreased N1 amplitudes compared to controls (Two-way ANOVA, p=0.0607, exploratory statistics: Student t-test, p=0.02, 0.004 for 0.1 and 10 cd*s/m^2^, respectively). The amplitude of P2 was not significantly different in rTg4510 compared to control mice at any of the ages tested (Fig. 2C). Furthermore, N1 and P2 latencies relative to the onset of the stimulus were not significantly different between genotypes (Fig. S4).

### Abnormalities in visual evoked potential responses correlates with tau pathology in young rTg4510

We characterized the progression of tau pathology in rTg4510 mice by biochemical tau analysis on brain homogenates and examined for phosphorylated tau species at the pS396 epitope. In rTg4510 mice at all ages, 55 kDa tau species, representing the human 4R0N transgenic P301L tau phosphorylated at the pS396 epitope were observed in the soluble fraction S1s by Western blotting. Hyperphosphorylated 4R0N tau-P301L appears as 64 kDa kDa bands in the soluble pellet fraction S1p as oligomeric tau and are regarded as the biochemical pre-tangle tau equivalent while the appearance of the 64 kDa bands in the sarkosyl-insoluble fraction P3 as aggregated tau is regarded as the biochemical equivalent to tangles detected by immunohistochemistry [30] (Fig. 3A, B).

We confirmed previous findings [22] that significant accumulation of hyperphosphorylated 64 kDa tau in soluble pellet S1p and insoluble P3 fraction was observed from 6 months of age in rTg4510 mice (One-way ANOVA, p<0.0001 for 3 vs. 6 months in both S1p and P3 fractions; Fig. 3D). In line with this accumulation of pathological tau species we confirmed the decrease of normal 55 kDa tau species. At 3 months of age, hyperphosphorylated 64 kDa was already detectable in both in S1p and P3 fractions confirming previous findings that oligomeric and aggregated tau species were present at this young age in rTg4510 mice. (Fig. 3C, D).

In order to evaluate whether the observed electrophysiological phenotype was associated to pathological tau levels, a correlation between N1 component of the VEP recorded during anesthesia and 64 kDa hyperphosphorylated tau species from S1p and P3 fractions was tested. We observed a significant positive correlation between both oligomeric and aggregated tau species and amplitudes of the N1 component in 3-6 month old rTg4510 (Simple linear regression, S1p: p=0.0196 and P3: p=0.0451; Fig. 3E). Namely, increased hyperphosphorylated tau species were associated with enhanced N1 amplitudes of the VEP response. On the other hand, late-stage rTg4510 mice (9-16 month old), did not show significant correlation between hyperphosphorylated tau levels and N1 of the VEP (Fig. 3F).

Next, we characterized the progression of tau pathology at the cellular level in the visual cortex and hippocampus by histology using AT8-immunoreactivity. Brain tissues from 3, 6, 9 and 16 month old rTg4510 and control animals were probed with the AT8 antibody. No AT8+ neurons were found in the brains from control animals at any age tested (Fig. 4A, B). From 3 months and older, tau phosphorylation at pS202/T205 increased and accumulated in the cell soma throughout the cortical layers and the hippocampus. In 3 month old rTg4510, AT8 immunoreactivity was mostly abundant in cortical layer II/III, while the distribution was more widespread throughout all layers of the visual cortex in 6, 9 and 16 month old rTg4510 mice. In addition, the AT8+ immunoreactivity was present in neuronal soma and processes in 3 months rTg4510 mice, and in 6-16 month old mice, the AT8 staining was restricted to the soma. There was a significant effect of age on AT8+ immunoreactivity in the visual cortex in rTg4510 mice (One-way ANOVA, p<0.0001; Fig. 4C). Normalized AT8+ cell counts in visual cortex of 3-9 months were similar and significantly increased at 16 months of age (One-way ANOVA, p=0.0039, <0.0001, for 6 vs. 9 and 9 vs. 16 months). In the CA1 area of the ventral hippocampus, a different progression was observed. At 6 months, rTg4510 displayed significantly more tau phosphorylation compared to 3 month old animals (One-way ANOVA, p<0.0001). The highest tau phosphorylation was quantified at 9 months of age while in 16 month old rTg4510 mice, the CA1 pyramidal layer was absent due to extensive neurodegeneration at this age. The AT8 immunoreactivity in CA1 is consistent with findings in the S1p fraction of the lysates containing both cortex and hippocampus by Western Blot. In visual cortex we observed a consistent decline in cortical thickness from 6 months of age onwards (One-way ANOVA, p<0.0001; Fig. 4D, E), suggesting gradual neuronal loss with increasing tau pathology. While neurodegeneration in the CA1 occurs from approximately 12 months [22], our data indicates that neurodegeneration in the visual cortex begins earlier, which may potentially explain the discrepancy between pre-tangle tau quantified in S1p by Western Blot and AT8 immunostaining.

**Figure 4.**
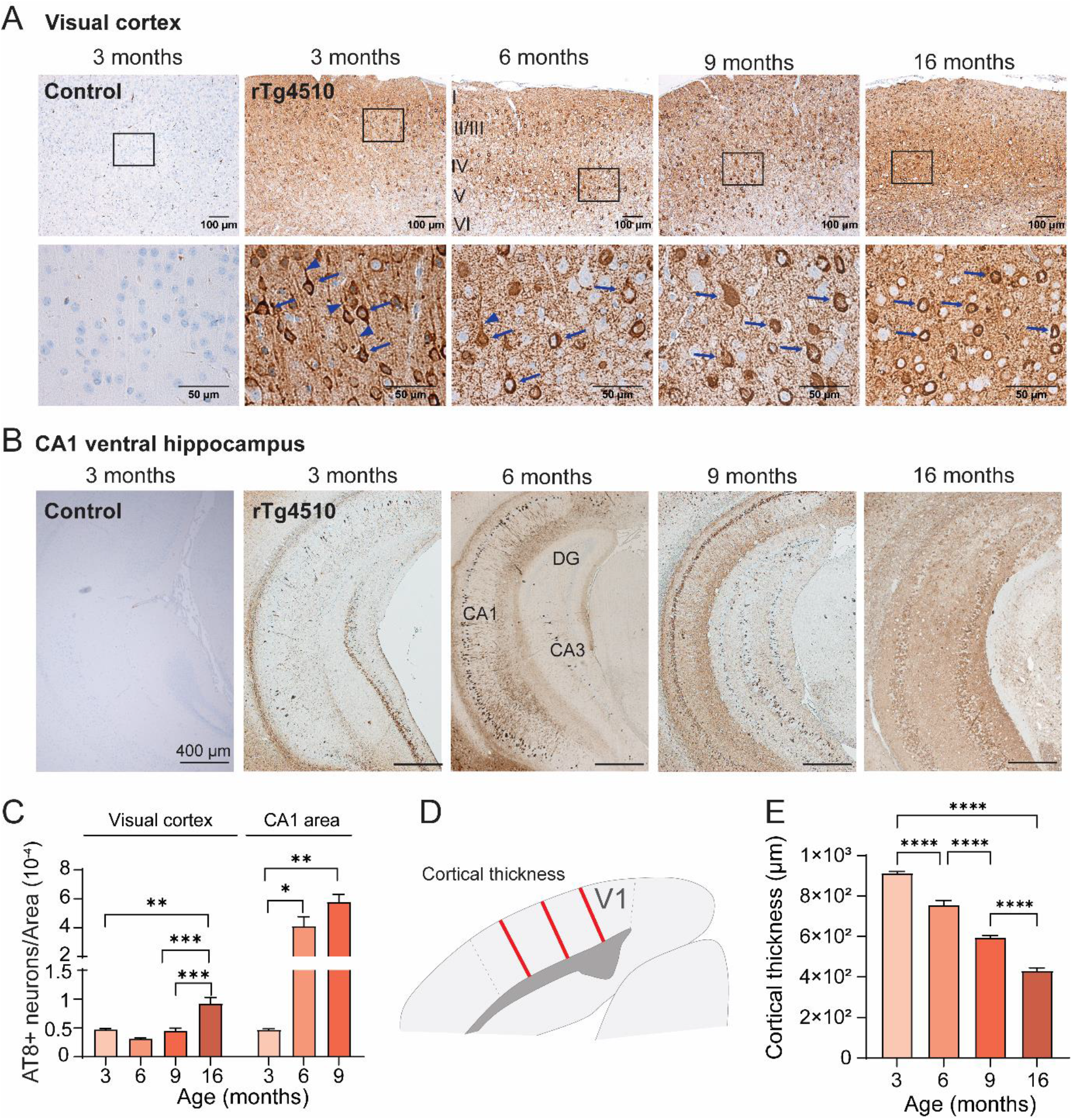
Pathological progression of tau hyperphosphorylation in the visual cortex and ventral hippocampus of rTg4510 mice. (A) Exemplary images of visual cortex and (B) ventral hippocampus of 3 month old control and 3, 6, 9 and 16 month old rTg4510 mice. AT8 antibody was used to evaluate phosphorylated tau localization in the tissue and visualized with horseradish peroxidase conjugated secondary antibody. High magnification images of (A) indicate the different localization of the AT8 staining to different neuronal compartments in the visual cortex. Arrows indicate soma and arrow-heads indicate processes. Note presence of AT8+ staining in processes only in 3 month old rTg4510. High magnification image of 3 month old rTg4510 is from layer II/III and of 6-16 month old mice from layer IV/V. (C) AT8 positive cells in rTg4510 mice visual cortex area and CA1 area of the ventral hippocampus and normalized to the size of the area of counting. AT8+ cells for each age group were quantified from four levels spaced out by approximately 40 μm (n=2-4). (D) Schematic of measurements of cortical thickness. (E) Cortical thickness of rTg4510 mice at 3, 6, 9 and 16 months of age (n=3-4). One-way ANOVA, effect (*) p<0.05, (**) p<0.01, (***) p<0.001, (****) p<0.0001. Data are shown as mean ± SEM. DG – dentate gyrus

Taken together, biochemical and histological analysis of progression of tau pathology indicated that abnormalities in visual function associated with early tau pathology and preceded neurodegeneration.

### Enhanced amplitude of N1 was confirmed in awake freely moving rTg4510 mice

Due to the potential of anesthesia to interfere with the cortical responses, we investigated whether the phenotype was preserved in awake mice. To this end, we recorded VEP responses in 6 month old awake freely moving mice (Fig. 5A). Overall, and independently of the genotype, N1 amplitude was lower in awake freely moving mice compared to anesthetized (Two-way ANOVA, p<0.0001; Fig 3B). On the other hand, the P2 component was enhanced in awake compared to anesthetized mice (Two-way ANOVA, p<0.0001). Additionally, both N1 and P2 latency was shortened in awake compared to anesthetized mice (Two-way ANOVA, p<0.0001 for N1 and P2; Fig. 5C).

**Figure 5.**
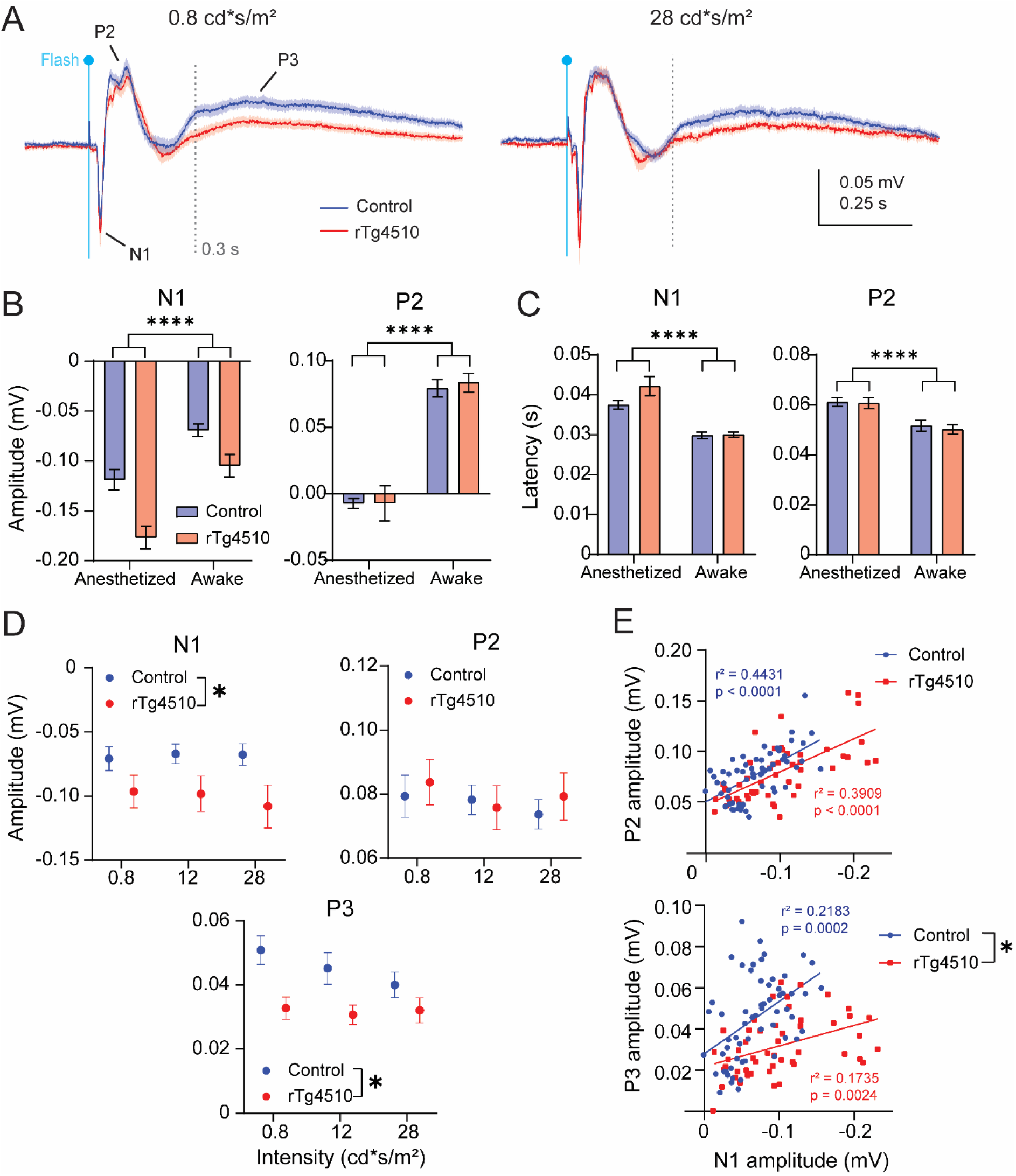
VEP recordings in 6 month old awake and freely moving control and rTg4510 mice. (A) Average VEP waveforms at lowest (left) and highest (right) tested intensity (0.8 and 28 cd*s/m^2^, respectively) in control (blue, n=20) and rTg4510 (red, n=15) animals. (B) Comparison of N1 (left) and P2 (right) amplitudes and (C) latencies between VEP recordings obtained from anaesthetized (1 cd*s/m^2^; n=17 and 23 for control and rTg4510, respectively) and awake mice (0.8 cd*s/m^2^; n=20 and 15 for control and rTg4510 mice, respectively). Two-way ANOVA, (****) p<0.0001. (D) Amplitudes of N1, P2, and P3 components of VEP recordings in control (blue, n=20) and rTg4510 (red, n=15) animals. Two-way ANOVA, (*) p<0.05. (E) Correlation of N1 and P2 (top) and N1 and P3 (bottom) between control and rTg4510 animals. Simple linear regression within genotype and student t-test, genotype effect, (*) p<0.05. Data are shown as mean ± SEM.

The modulation of the N1 and P2 amplitude elicited by incremental light intensities observed in anesthetized mice was absent in awake mice for the range of intensities tested (Fig. S5). Our recording settings in awake mice also allowed to visualize the late P3 component, which was not investigated in anesthetized mice. The amplitude of the P3 was significantly decreased in rTg4510 compared to control animals (Two-way ANOVA, p=0.0187; Fig 5D). Importantly, the amplitude of N1 was significantly increased in awake rTg4510 mice compared to controls (Two-way ANOVA, p=0.0447; Fig. 5D), in agreement with our observations in anesthetized mice. In line with recordings in anesthetized mice, P2 amplitude was not significantly different between genotypes (Fig 5D). Latencies of all components analyzed were not altered in rTg4510 mice compared to controls (Fig. S6).

A correlation analysis was carried out between amplitudes of N1 and P2, and N1 and P3, to evaluate potential interdependencies (Fig. 5E). There was a significant positive correlation between N1 and P2 for control and rTg4510 mice, indicating that enhancement in N1 amplitude resulted in higher P2 amplitude (Simple linear regression, p<0.0001 for both control and rTg4510 mice; Fig. 5E). A significant positive correlation was also found between N1 and P3 components in both genotypes (Simple linear regression, p=0.0002 and 0.0024 for control and rTg4510, respectively). However, the correlation observed in rTg4510 was significantly lower compared to control mice, with increasing N1 amplitude resulting in lower P3 amplitude in rTg4510 (Simple linear regression, p=0.0024; Fig. 5E). This loss of correlation suggests that in 6 months old rTg4510, mechanisms underlying the N1 and P3 components are altered.

In summary, we found a notable influence of anesthesia on VEP waveforms. Nevertheless, in both the anesthetized and awake paradigm, we detected an enhancement in N1 amplitude in 6 month old rTg4510 mice compared to controls.

## Discussion

The aim of our study was to investigate whether neuronal functioning in the visual pathway reflects early- and late-stage changes associated with progression of tau pathology could be measured using electrophysiology. We found marginal alterations in ERG responses within age groups in rTg4510 mice despite the presence of hyperphosphorylated pre-tangle tau in the photoR and INL layers. However, a decline in b-wave amplitude was observed as a function of age in transgenic mice. Additionally, we discovered an attenuation of the PhNR response from 6 months of age indicative of functional impairment of RGCs where hyperphosphorylated pre-tangle tau was also detected. Cortical visual evoked potentials were enhanced in young rTg4510 mice, whereas a decrease was observed in older animals likely due to neurodegeneration as indicated by decreased cortical thickness. Levels of oligomeric and aggregated hyperphosphorylated tau species were positively correlated with changes in the cortical evoked potentials in young rTg4510.

The steady reduction in cortical thickness from 6 months of age and presence of AT8 immunoreactive tau species from 3 months of age, suggest that the increase in amplitude of the cortical evoked waveform in young animals precedes neurodegeneration induced by tau pathology. The early neuronal manifestations of the effects of human tau overexpression in the visual pathway in rTg4510 mice, prior to reported neuronal loss and cognitive defects [21,33], indicates that the assessment of visual processing may represent a useful biomarker also for early clinical diagnosis.

We first investigated the consequences of tauopathy in visual processing by recording retinal responses elicited by a flash of light. Overall, we found that a- and b-wave amplitudes and latencies were unchanged between the two genotypes within age groups despite detectable hyperphosphorylated tau in photoreceptor layer of rTg4510 animals already from 3 months of age. Similarly to this study, unchanged scotopic ERG responses were reported in a hTau mouse model at 5 and 17 months despite evident AT8 immunoreactive hyperphosphorylated tau species in the INL and RGC layers [16]. The observed age-dependent decline of the b-wave in rTg4510 might be attributed to hyperphosphorylated tau in the INL and abnormal aging of this structure [14]. As a-wave denotes photoreceptor activation while b-wave represents INL responses [34,35], we conclude that function of these neuronal populations remain intact at early stages of tau pathology but undergo age-dependent alterations.

RGC function was specifically investigated using a PhNR protocol [36]. A significant reduction in PhNR responses was found in rTg4510 animals from 6 months of age even though AT8 immunoreactivity was consistently detected in RGC layers of rTg4510 retinas starting from 3 months of age. The decline in RGC evoked responses at older ages is consistent with observations in glaucoma models, which are characterized by degeneration of the optic nerve and loss of retinal ganglion cells [37,38]. Further, accumulation of hyperphosphorylated tau in the retina has been associated with a reduction of RGC density and optic nerve thinning in rTg4510 mice [14]. Taken together, our findings show that impairments in retinal visual processing were absent in young rTg4510 and developed in older ages possibly as a consequence of hyperphosphorylated tau-dependent retinal degeneration.

Alongside retinal processing, we evaluated the effects of tau overexpression on the function of the visual cortex as the final relay in the visual sensory pathway. Analysis of cortical responses to a flash of light revealed that N1 amplitudes were augmented from 3 to 9 months of age in rTg4510 mice compared to controls. VEP waveform architecture was significantly altered by anesthesia, with increased N1 amplitude and decreased P2 amplitude, and a prolonged latency of both components. However, despite the notable influence of anesthesia on VEP waveforms, genotype differences in N1 amplitudes were detectable in both awake and anesthetized animals. While trends towards enhanced N1 amplitudes in rTg4510 mice were recently reported in visual cortex at 5 and 8 months, after visual stimulation [17], to our knowledge this is the first study reporting significant early changes in visual evoked potentials as a consequence of tau pathology.

Multiple mechanisms might contribute to the observed selective alterations in cortical VEPs in rTg4510 animals, rather than local defects at the level of the eye measured by ERGs. Visual cortical neurons are the source of VEPs as they receive feed-forward thalamic inputs in the sensory pathway and feed-back signals on the cortico-cortical axis [39–42]. As subcortical areas such as thalamus do not express human tau in rTg4510 mice, a possible impairment of the visual cortical circuits at single cell level likely occurs predominantly in visual cortex [21]. Aberrant excitability of cortical pyramidal neurons and inhibitory interneurons as well as their synaptic interconnections might reshape the observed visual processing in early stages of tau pathology [39,42,43]. A potential impairment of the excitability of pyramidal neurons could be responsible for the visual processing alterations at early stage as found in prefrontal cortex and CA1 hippocampal layer in older rTg4510 mice [24,44,45]. A decline in firing rates and increase in silenced cortical pyramidal neurons was reported in young rTg4510 [46,47]. Moreover, hippocampal recordings from rTg4510 animals show decline in pyramidal neuron firing rates [27], reduced excitability [48,49] as well as depletion of pyramidal neurons in the CA1 region all prior to severe neurodegeneration [22,49]. Secondly, a dysfunction of GABAergic transmission could explain VEP alterations. Indeed, THIP, a GABAAR agonist, depressed N1 and P2 amplitudes and augmented P3 component of VEP response [50], while bicuculline, a GABA_A_R antagonist, enhanced N1 amplitudes in somatosensory cortex of naïve mice [51]. Additionally, PET imaging revealed reduced uptake of GABAergic [^11^C]flumazenil tracer in young rTg4510 animals [25]. Finally, abnormal synaptic connectivity may influence cortical processing as disruption in axonal transport caused by hyperphosphorylated tau-triggered microtubule destabilization found *in vitro* might attenuate local activity as well as long-range neuronal connectivity in hippocampus and cortex [4,25,27,46]. The decrease in correlation between VEP peaks (N1 versus P3) may indicate a general alteration in single-cell neuronal synchrony and synaptic transmission amongst pyramidal cells and inhibitory interneurons in visual cortex as a result of hyperphosphorylated tau accumulation [21]. The present findings do not provide understanding of the precise mechanism(s) responsible for alterations in VEP response in young rTg4510 mice. In this respect, further studies aimed at probing single cell activity, for example using calcium imaging of identified neuronal populations, will be crucial.

To investigate whether the early alterations in visual processing in rTg4510 mice are associated with tau pathology, we quantified the levels of hyperphosphorylated tau in the brain and evaluated their link to VEP responses across ages. The 64 kDa hyperphosphorylated tau species resemble part of the tau pathology found in AD brains [3]. Consistently with previous studies, we already detected levels of 64 kDa hyperphosphorylated tau species in 3 month old brains of rTg4510 mice [22]. AT8 immunoreactivity was similar in the visual cortex and hippocampus in 3 month old rTg4510. However, in older animals levels of pathological tau were higher in the ventral hippocampal area compared to visual cortex as indicated by a more severe increase of AT8 immunoreactivity in this anatomical region. The presence of AT8+ neurons in the visual cortex of rTg4510 was associated with gradual decline of cortical thickness from 6 months of age, suggesting that neuronal loss occurs earlier in visual cortex compared to the CA1 of hippocampus [22]. Finally, our data show that N1 amplitudes were negatively correlated with oligomeric and aggregated hyperphosphorylated tau species in young animals, and with the pre-tangle tau in the visual cortex, indicating that the early change in VEP response was dependent on pathological tau levels.

In summary, our findings show early consequences of human tau overexpression in the visual pathway in the rTg4510 mouse tauopathy model. We found a novel age- and pathology-dependent VEP biomarker prior to severe neuronal loss. Interestingly, the neuronal signature observed in VEP recordings is associated with hyperphosphorylated tau exclusively in young rTg4510 animals. Further experiments aiming at understanding the precise mechanisms underlying impairments in visual processing as a result of early tau pathology are warranted. More importantly, our findings support the mounting evidence that VEPs could serve as biomarkers for early stages of neurodegenerative disorders [52–54], as also suggested for other sensory domains [55,56]. While the present study indicates its potential usefulness as a diagnostic tool of early tauopathy, visual processing evaluation could also represent a valuable target engagement biomarker for tau reducing therapeutic principles.

## Acknowledgements

This work was supported by the Innovation Fund Denmark (Ref. no. 9065-00019B). We are grateful for the technical assistance of Lise Maj Schrøder-Hansen, Kirsten Christensen, Anette Bredal Christiansen and Annette Bjørn at H. Lundbeck.

## Supplementary materials

**Fig. S1.**
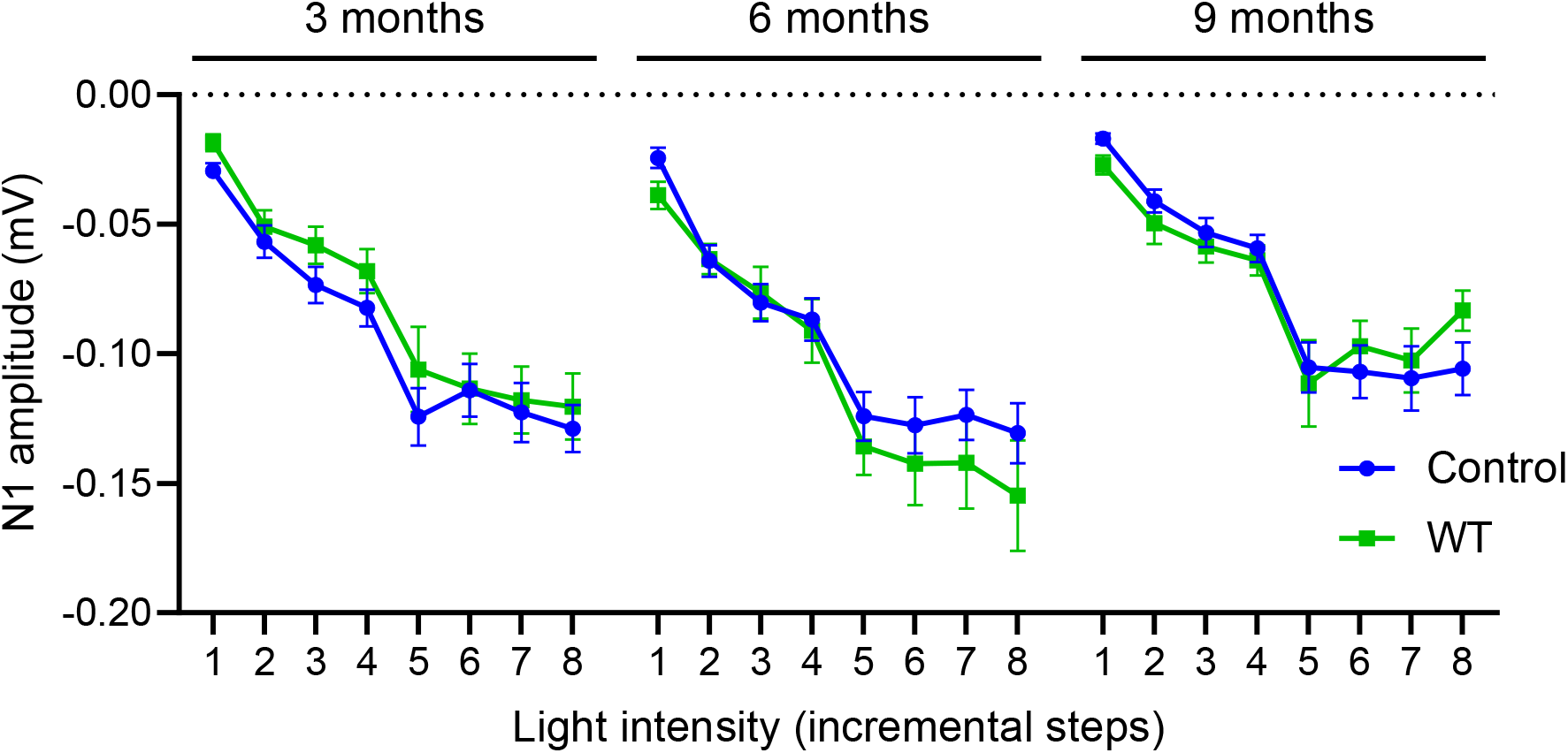
N1 amplitudes of VEP recorded from anesthetized tTA (control, blue, n=20-22) and non-transgenic mice (WT, green, n=7-10) at 3, 6 and 9 months of age. X-axis represents stimulation intensities (1-8 for 0.003, 0.001, 0.03, 0.01, 0.3, 0.1, 1, 10 cd*sm^2^, respectively). Data are shown as mean ± SEM.

**Fig. S2.**
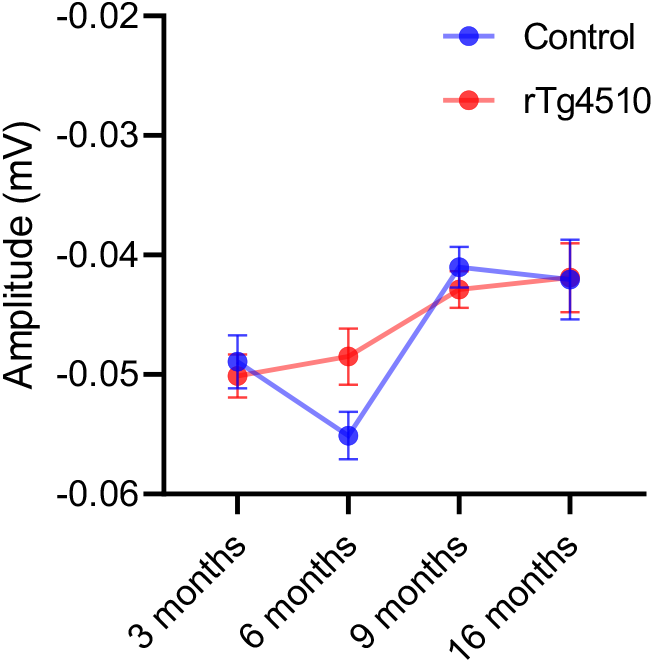
Amplitudes of the a-wave of the ERG as a function of age in tTA (control, blue, n=15-22) and rTg4510 (red, n=19-24) mice. Data are shown as mean ± SEM.

**Fig. S3.**
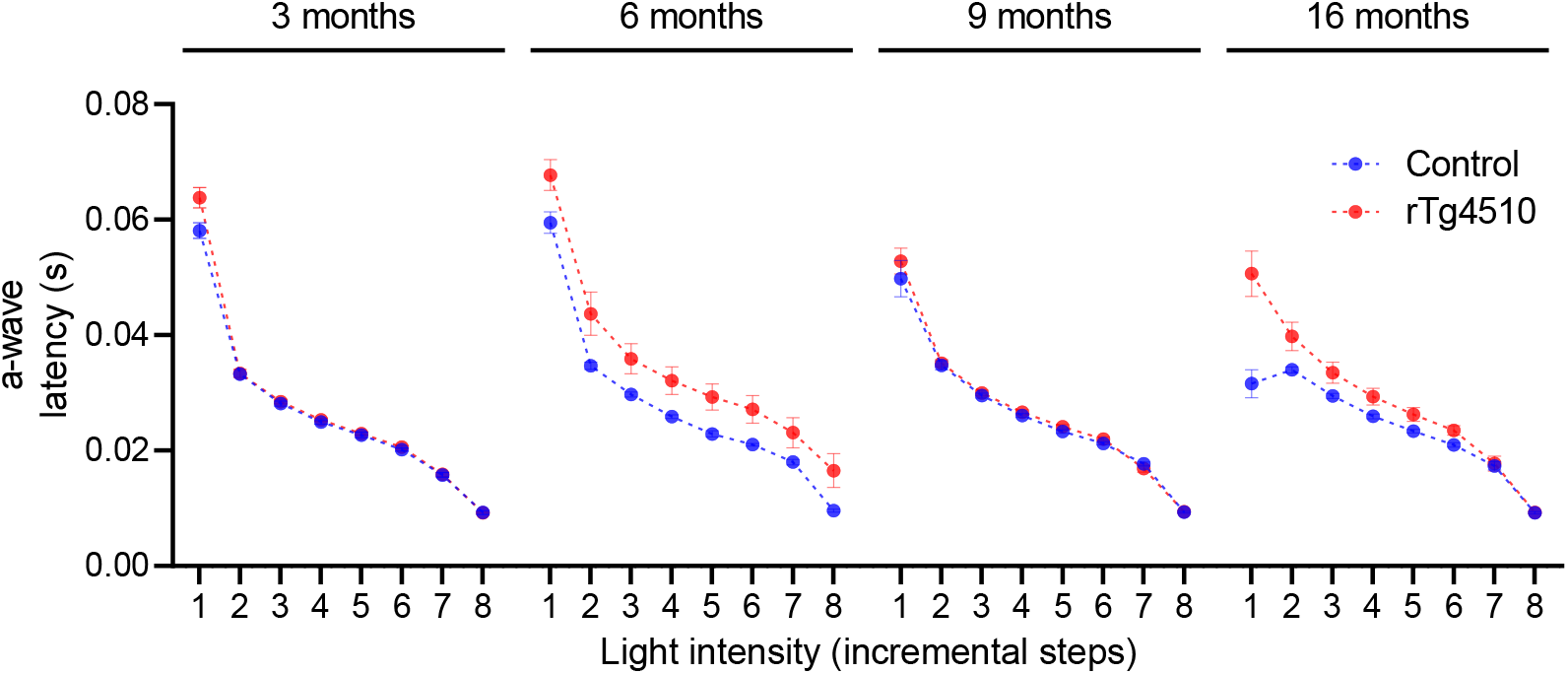
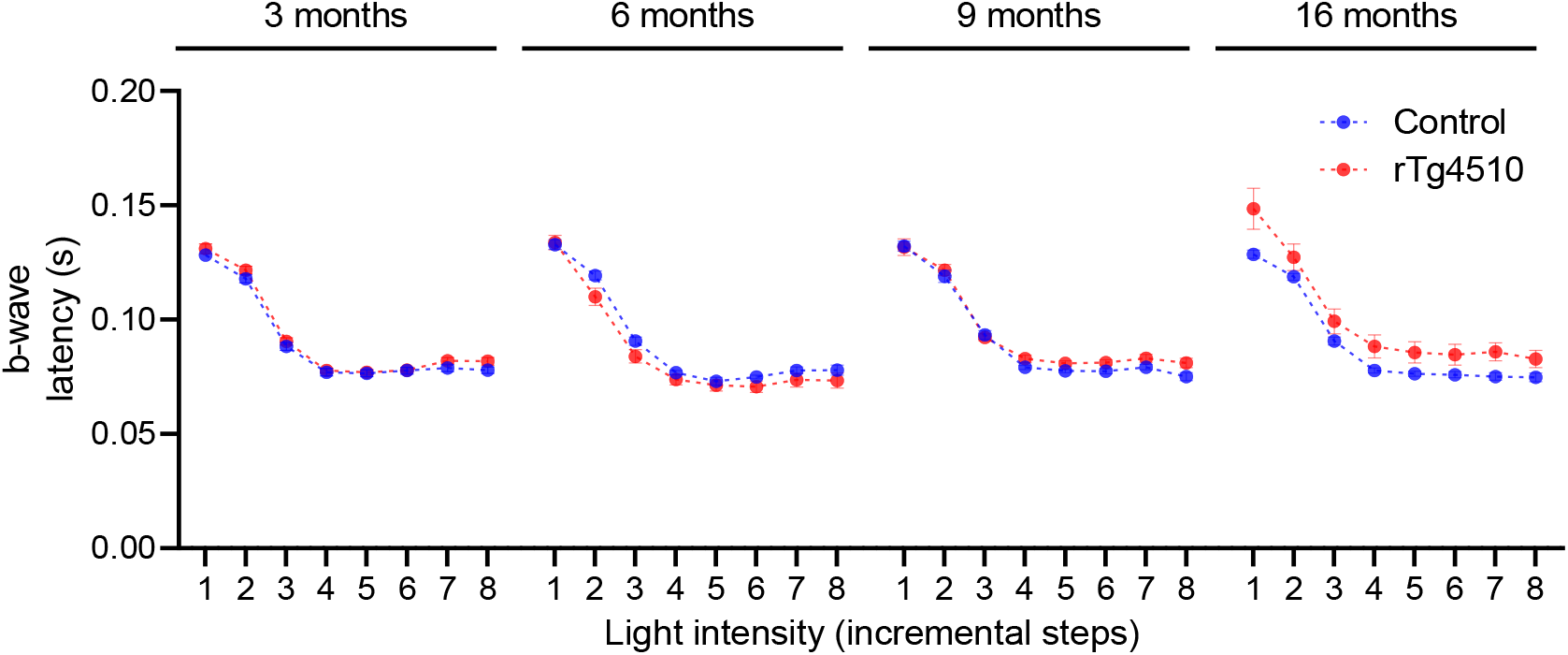
Latencies of a- (top) and b-wave (bottom) of the ERG responses in tTA (control, blue, n=15-22) and rTg4510 (red, n=19-24) mice at 3, 6, 9 and 16 months of age. X-axis represents stimulation intensities (1-8 for 0.003, 0.001, 0.03, 0.01, 0.3, 0.1, 1, 10 cd*sm^2^, respectively). Data are shown as mean ± SEM.

**Fig. S4.**
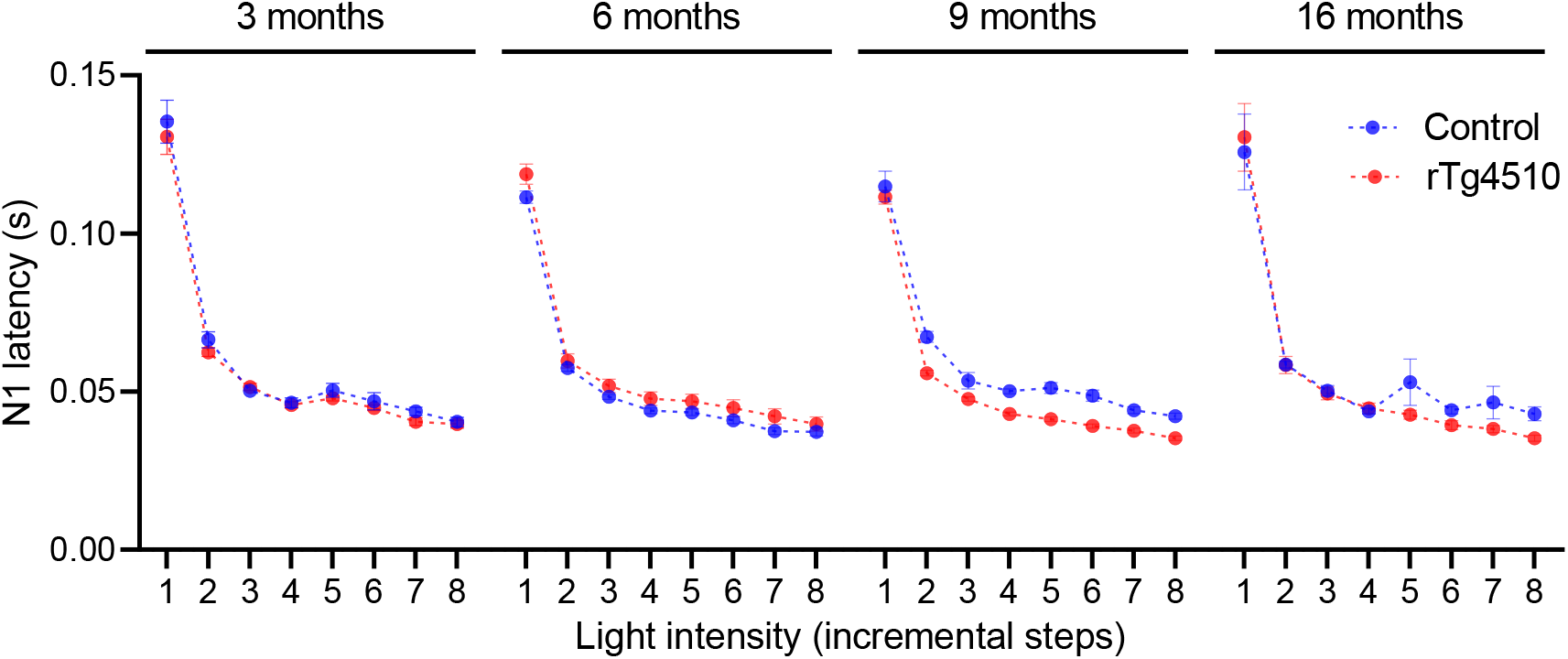
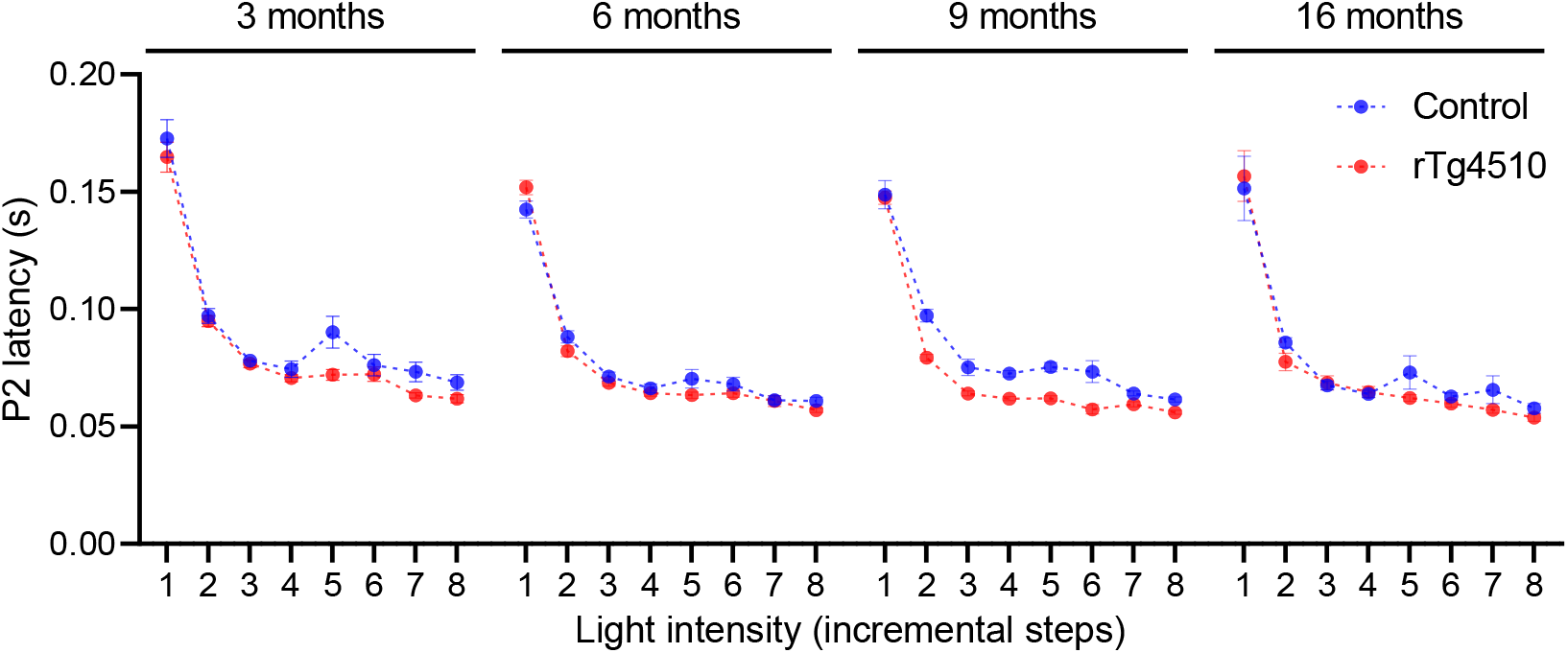
Latencies of N1 (top) and P2 (bottom) of the VEP responses in anesthetized tTA (control, blue, n=15-22) and rTg4510 (red, n=19-24) mice at 3, 6, 9, and 16 months of age. X-axis represents stimulation intensities (1-8 for 0.003, 0.001, 0.03, 0.01, 0.3, 0.1, 1, 10 cd*sm^2^, respectively). Data are shown as mean ± SEM.

**Fig. S5.**
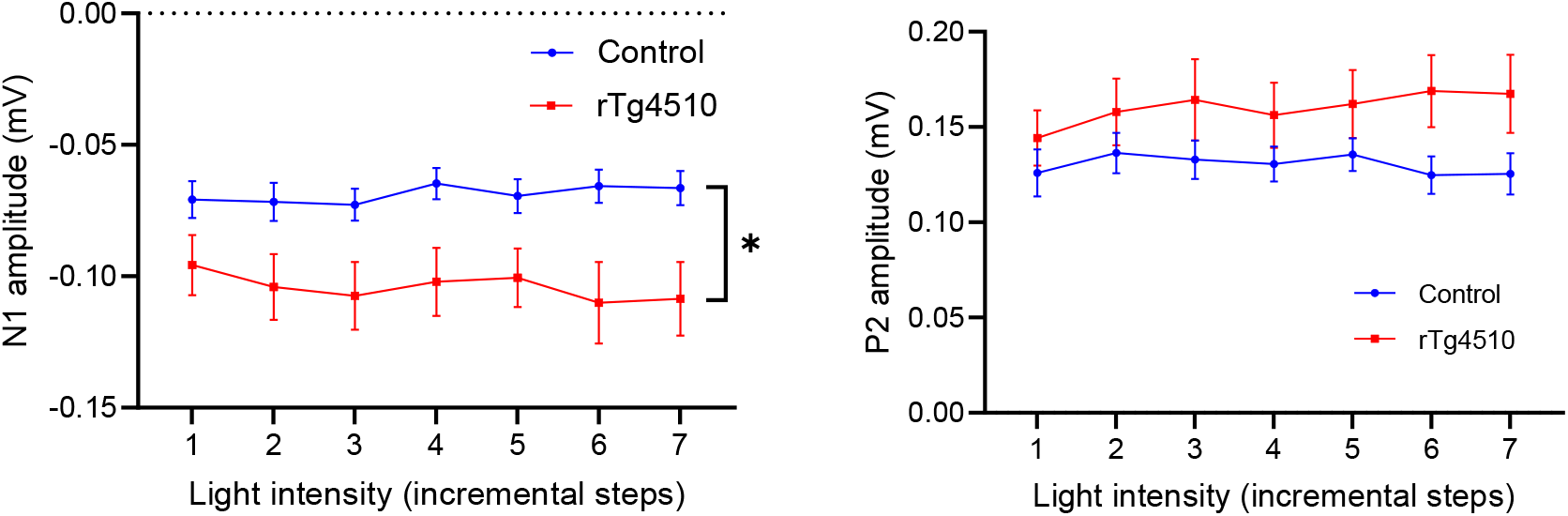
Amplitudes of N1 (left) and P2 (right) of the VEP recordings in 6 month old awake freely moving tTA (controls) and rTg4510 (n=20 and 15, respectively) at incremental intensities (1-7 on the x-axis corresponds with 0.8, 3.44, 7.2, 11.8, 17.3, 22.4, 28.3 cd*s/m^2^, respectively). Two-way ANOVA, (*) p<0.05. Data are shown as mean ± SEM.

**Fig. S6.**
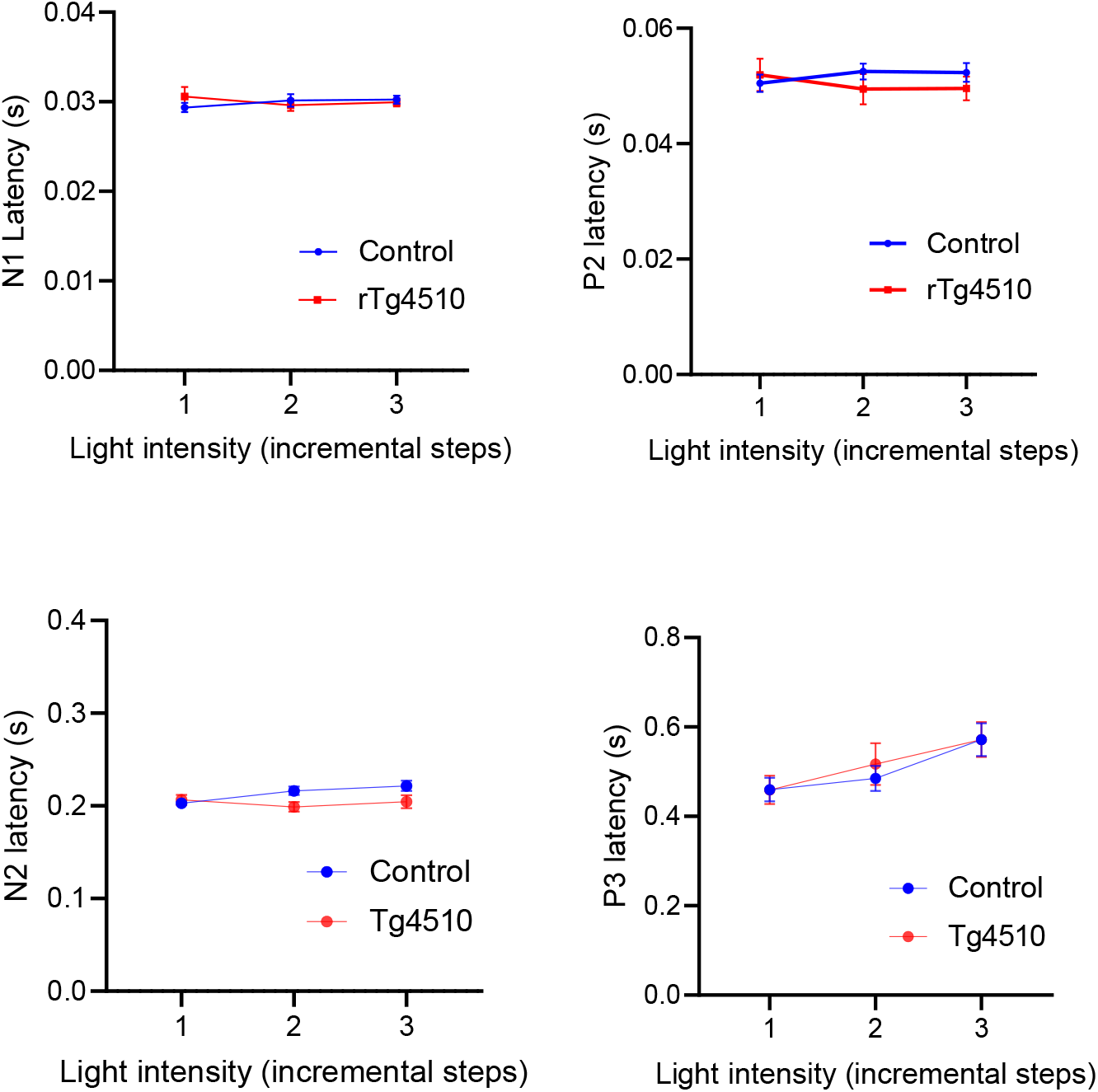
Latencies of N1, P2, N2 and P3 of the VEP recordings in 6 month old awake freely moving tTA (controls, blue, n=20) and rTg4510 (red, n= 15) at incremental intensities (1-3 on the x-axis corresponds with 0.8, 11.8, 28.3 cd*s/m^2^, respectively). Data are shown as mean ± SEM.

